# Common genetic variants and health outcomes appear geographically structured in the UK Biobank sample: Old concerns returning and their implications

**DOI:** 10.1101/294876

**Authors:** Simon Haworth, Ruth Mitchell, Laura Corbin, Kaitlin H Wade, Tom Dudding, Ashley Budu-Aggrey, David Carslake, Gibran Hemani, Lavinia Paternoster, George Davey Smith, Neil Davies, Dan Lawson, Nicholas Timpson

## Abstract

The inclusion of genetic data in large studies has enabled the discovery of genetic contributions to complex traits and their application in applied analyses including those using genetic risk scores (GRS) for the prediction of phenotypic variance. If genotypes show structure by location and coincident structure exists for the trait of interest, analyses can be biased. Having illustrated structure in an apparently homogeneous collection, we aimed to a) test for geographical stratification of genotypes in UK Biobank and b) assess whether stratification might induce bias in genetic association analysis.

We found that single genetic variants are associated with birth location within UK Biobank and that geographic structure in genetic data could not be accounted for using routine adjustment for study centre and principal components (PCs) derived from genotype data. We found that GRS for complex traits do appear geographically structured and analysis using GRS can yield biased associations. We discuss the likely origins of these observations and potential implications for analysis within large-scale population based genetic studies.

## Main

Many recent and ongoing research programmes aim to systematically identify genetic contributions to complex traits and undertake applied epidemiological analyses using genotype data. Irrespective of source, latent structure within a dataset can be very important when performing these analysis, as structural alignment between ancestry and genotypes, health outcomes and geography has potential to induce artefactual relationships^1^. Current methods to account for structure include proxy measurement and adjustment for latent structure within datasets (mainly using PCs or measures of actual geographic location^2–4^).

Recent developments in resources, applications and understanding warrant a re-exploration of latent structure in datasets. Prior to 2015, very large samples were only achieved by aggregation of smaller studies whose structural properties and geographical footprints were neither detectable within single studies nor coordinated across the collection of studies. Now analysis can be undertaken in very large individual collections with the capacity to capture a single geographical footprint, such as UK Biobank^5^. With increased sample size and statistical power, there is now potential to discover a broader range of genetic effects that might conceivably capture characteristics of the structural properties or geographical footprint of the dataset. This sits in the context of a growing appreciation of fine-scale population structure within the British population^6^.

These changing circumstances are relevant for applied epidemiological analyses which have developed substantially with their exploitation of reliable genetic association results. A good example of this is Mendelian randomization, which aims to escape confounding in observational associations by using genetic variation to proxy risk factors of interest^7^. Recent literature has focused on maximising the use of the current wave of genetic association evidence and accounting for undesirable pleiotropic effects of single variants^8^. This activity, however, has largely assumed that structure is addressed during the discovery of associated genetic variants. Under-appreciated structure in genetic datasets challenges the assumption that genetic instruments are not related to potentially confounding features^9^.

As an exemplar, we examined whether there is previously under-appreciated structure in a well understood, ethnically and geographically homogenous resource. In the Avon Longitudinal Study of Parents and Children (ALSPAC)^10,11^, we studied mothers who were recruited during pregnancy in the Bristol area (South West UK) in the early 1990s. We undertook chromosome painting^12^ to describe fine-scale relatedness between each mother and each of the regions of the Peopling of the British Isles (PoBI) project^6^. We summarised each mother’s ancestral lineage as a mixture of the PoBI regions, allowing us to estimate the educational attainment that those regions would have, were the ALSPAC mothers’ education levels explained by this variation. In doing this a pattern for lower educational attainment in lineages originating from the regions immediately surrounding Bristol (Figure 1) and higher educational attainment in more geographically distant lineages was observed. Distant lineages are likely only represented in ALSPAC by individuals or families who had migrated, and we anticipate that the educational attainment of people who migrate for economic reasons differs from people who do not. Educational attainment is therefore aligned to subtle genetic differences even in this apparently geographically and ethnically homogenous population and this is coincident with axes of ancestry.

**Figure 1:**
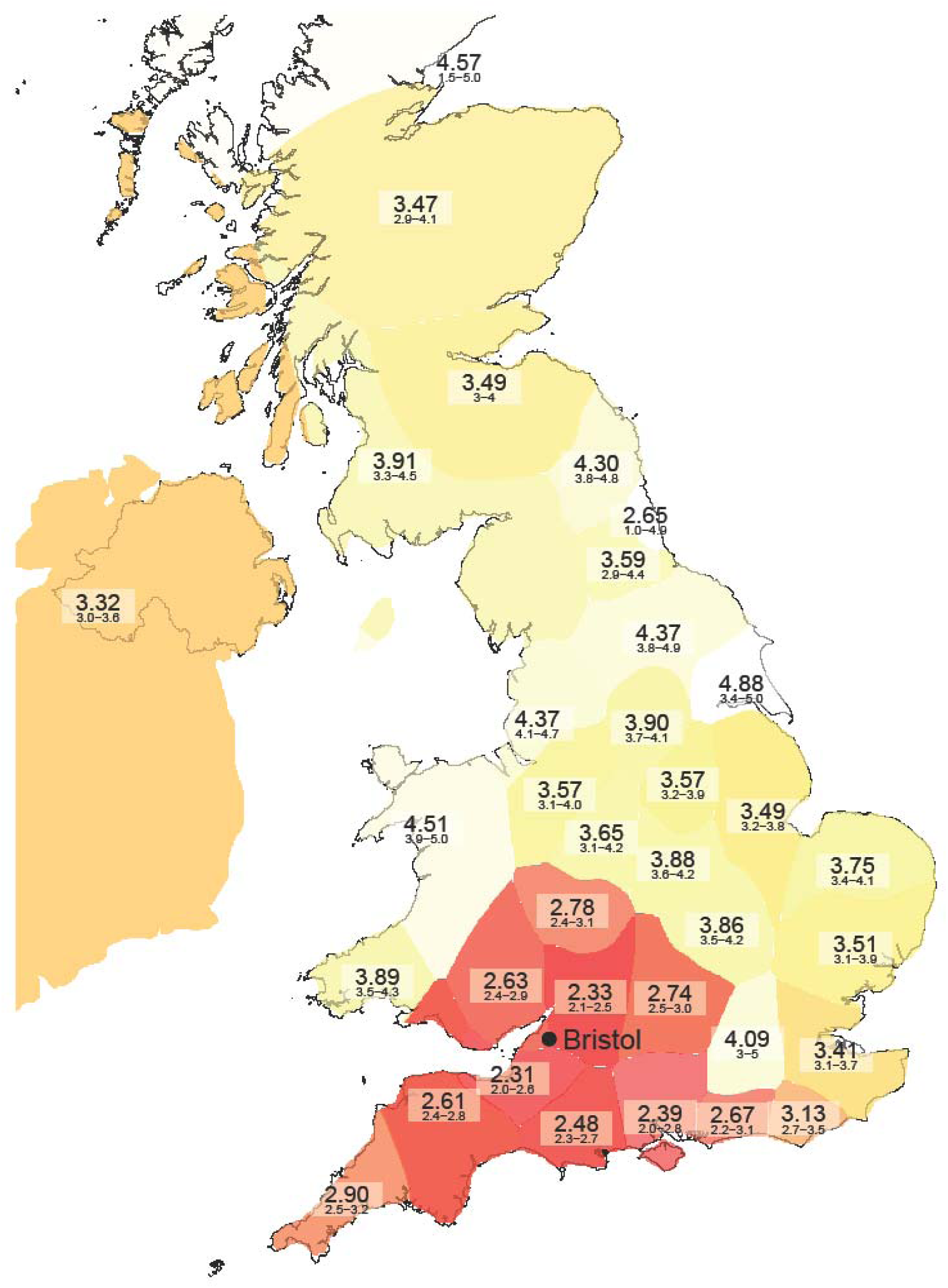
Within-UK ancestry predicts migration that confounds education: Estimated educational attainment of the UK, when seen only through the ALSPAC cohort based in Bristol. Scores are 1: Vocational, 2: CSEs, 3: O-levels, 4: A-levels, 5: degree. The predicted mean education for each region is given, along with 95% confidence intervals estimated by bootstrap resampling of individuals. Each region is coloured by predicted mean education. See online methods for details.

The structure in ALSPAC was detected here using a method which is highly sensitive to ancestry. With greater power, it is entirely possible the same phenomena may become detectable in more routine analytical procedures. We therefore turned to UK Biobank, an exceptional resource containing a catalogue of health, disease and genotype data of almost half a million participants^5,13^ Conceptually the UK Biobank is analogous to a super-imposition of multiple ALSPACs, each of which recruited participants living near a study assessment centre. This design gives UK Biobank the capacity to represent a broad spectrum of UK ancestry and structure, but is also sensitive to important sampling phenomena including self-selection. The hurdles of location and attendance (less than 6% of individuals contacted by UK Biobank chose to participate^14^) are likely to influence the nature of the resultant participant collection and are related to behaviours with heritable contributions^15^. This may create collider biases^16,17^ which have the ability to induce association between otherwise independent variables.

We examined whether genotypes are structured using genome-wide association studies (GWAS) for North/South and East/West axes of birth location (on a metre grid scale from an origin South West of the UK) using PLINK^18^. Analysis of genetic data was performed within individuals of white British ancestry with non-missing data on birth location (n=321,439). GWAS for birth location identified that single variants are associated with geography within UK Biobank. An unadjusted model produced distorted and inflated plots with evidence for association at variants across the autosome. After adjustment for genotyping array, 40 PCs and a factor variable representing UK Biobank assessment centre single variants remained associated with birth location (figure S1).

Rather than using single genetic variants, empirical epidemiological analyses often use genetic risk scores (GRS)^19,20^. As exemplars, we took genetic variants and weightings associated with educational attainment, height and body mass index (BMI) from published genome-wide meta-analyses^21–23^. Using an approach that is widespread in applied analyses, we derived weighted and unweighted GRS for the three traits based on variants with p<5e-08 and p<1e-05 in the discovery sample. We used general additive models^24^ in the ‘mgcv’ package (version 1.8)^25^ within R (version 3.3.1)^26^, to test for non-linear relationships between GRS and geographical terms. All GRS tested were associated with birth location in an unadjusted model and a model that adjusted only for genotyping array. These associations attenuated but were not extinguished in models incorporating adjustment for 40 PCs and study centre, especially for educational attainment and North location at birth, where statistical adjustment had little impact on the fitted geographical distribution of the GRS (figure 2, table 1).

**Figure 2:**
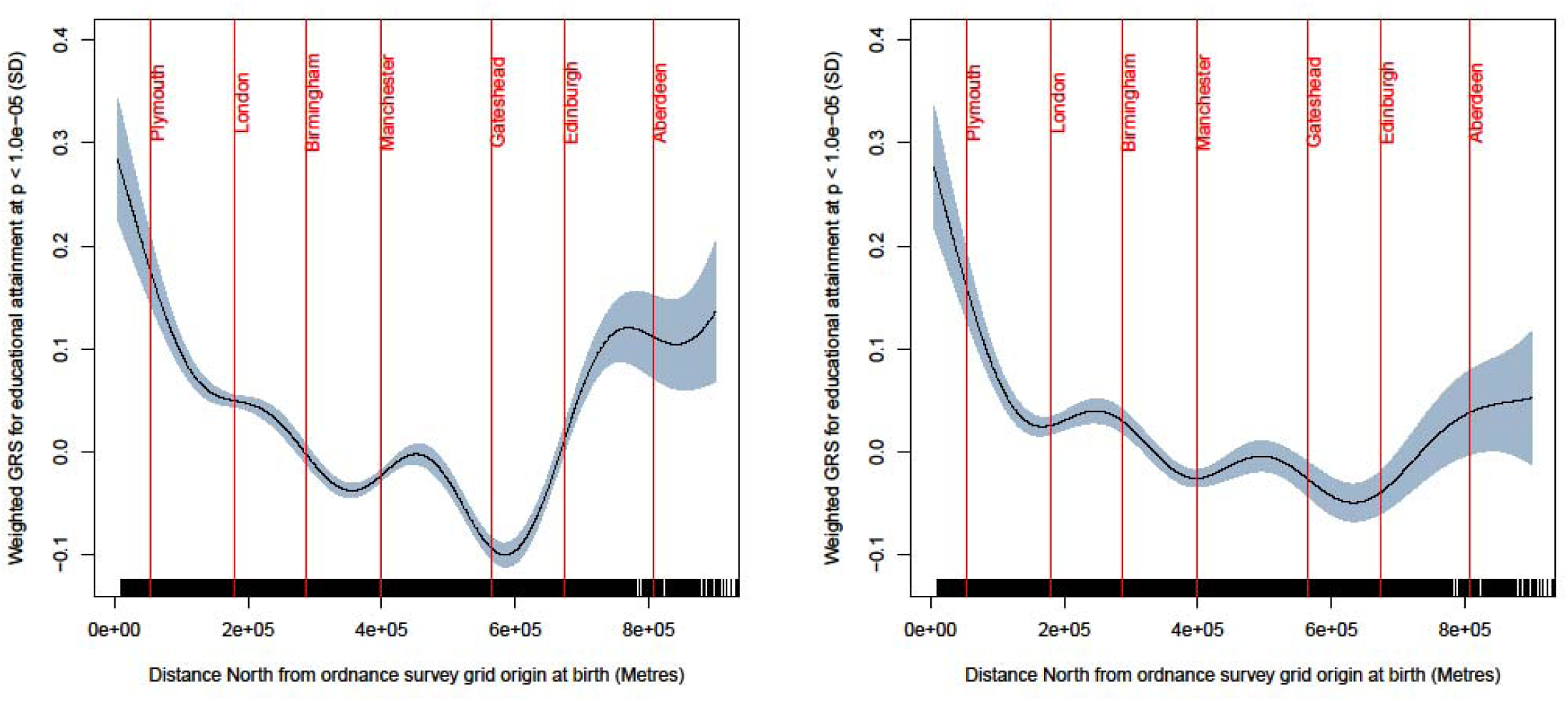
Fitted spline regression plots showing the non-linear distribution of GRS for educational attainment (weighted version, including variants with p<1.0e-05) in minimally adjusted model (left) and model after adjustment for 40 principal components and study centre (right). The centre of major population centres is marked for reference. The shaded area represents 95% confidence intervals

**Table 1.** Relationship between GRS and birth location within UK Biobank.

|  |  | P value for association between GRS and geographical term |  |  |  |  |  |  |  |
| --- | --- | --- | --- | --- | --- | --- | --- | --- | --- |
|  |  | Weighted GRS |  |  |  | Unweighted GRS |  |  |  |
|  |  | Model 1 | Model 2 | Model 3 | Model 4 | Model 1 | Model 2 | Model 3 | Model 4 |
|  |  | Educational attainment |  |  |  |  |  |  |  |
| P ( 5.0e-08) | North component | 2e-16 | <2e-16 | 6.4e-06 | 6.7e-06 | <2e-16 | <2e-16 | 1.3e-09 | 1.6e-06 |
|  | East component | <2e-16 | <2e-16 | 1.5e-09 | 6.0e-11 | <2e-16 | <2e-16 | 7.5e-14 | 1.3e-11 |
|  |  | Height |  |  |  |  |  |  |  |
|  | North component | <2e-16 | <2e-16 | 1.3e-05 | 0.14 | <2e-16 | <2e-16 | 4.6e-06 | 0.13 |
|  | East component | <2e-16 | <2e-16 | 0.00021 | 0.095 | <2e-16 | <2e-16 | 3.4e-05 | 0.046 |
|  |  | Body mass index |  |  |  |  |  |  |  |
|  | North component | 9.7e-07 | 9.9e-07 | 0.063 | 0.40 | 0.0013 | 0.0012 | 0.0032 | 0.58 |
|  | East component | 0.0036 | 0.0035 | 0.24 | 0.93 | 0.053 | 0.054 | 0.032 | 0.47 |
|  |  | Educational attainment |  |  |  |  |  |  |  |
| P ( 1.0e-05) | North component | <2e-16 | <2e-16 | <2e-16 | <2e-16 | 7.6e-11 | 8.5e-11 | 0.012 | 0.16 |
|  | East component | <2e-16 | <2e-16 | <2e-16 | <2e-16 | 9.7e-12 | 8.9e-12 | 0.0021 | 0.041 |
|  |  | Height |  |  |  |  |  |  |  |
|  | North component | <2e-16 | <2e-16 | 5.9e-05 | 0.16 | <2e-16 | <2e-16 | 0.00025 | 0.17 |
|  | East component | <2e-16 | <2e-16 | 0.00014 | 0.051 | <2e-16 | <2e-16 | 7.2e-05 | 0.014 |
|  |  | Body mass index |  |  |  |  |  |  |  |
|  | North component | 2.4e-09 | 2.5e-09 | 0.023 | 0.019 | 2.4e-10 | 2.6e-10 | 0.0029 | 0.074 |
|  | East component | 1.4e-13 | 1.7e-13 | 0.134 | 0.34 | <2e-16 | <2e-16 | 0.020 | 0.14 |

Table contents – p value for non-linear association between component of birth location and genetic risk score. For all models n=321,439. Statistical adjustment was performed as follows: model 1 – no adjustment; model 2 – adjustment for genotyping array only; model 3 – adjustment for genotyping array, 10 PCs and study participation centre; model 4 – adjustment for genotyping array, 40 PCs and study participation centre.

Having found evidence for association between genotypic variation and geography, we used general additive models to test for non-linear relationships between four exemplar complex traits and geography. Reported household income, measured BMI, reported age at completion of full time education and reported number of siblings showed strong evidence for geographical stratification (p<2e-16 for non-linear relationship between observed traits and axes of birth location).

We noted that structure in genotypes and phenotypes appeared geographically co-incident (example figure S2), which led us to explore the potential role of geography in confounding applied analysis. We tested for linear association between GRS and complex traits and examined whether the inclusion of non-linear terms for birth location as covariates altered the results, again using general additive models. These relationships changed in magnitude with the addition of non-linear terms for birth location (table 2), suggesting a role for residual confounding by geographical location. For example, the relationship between genetically predicted BMI and household income (pounds sterling per year per 1 standard deviation (SD) increase in GRS for BMI) changed from -335 in the unadjusted model to -251 (adjusted for 40 PCs and study location) to -229 (adjusted for 40 PCs study location and non-linear terms for birth location). Birth location captures neither the full extent of variation in fine ancestral structure (which predicts GRS) nor the full extent of geographically structured social and economic differences (which predict income). It is possible that these adjusted estimates therefore contain residual confounding and that the true impact of biases within this sample is larger than these results suggest.

**Table 2.** Linear relationships between observed traits and genetic risk scores in UK Biobank.

| Observed trait (unit per 1SD increase in GRS) | N | Weighted GRS |  |  |  | Unweighted GRS |  |  |  |
| --- | --- | --- | --- | --- | --- | --- | --- | --- | --- |
|  |  | Model 1 | Model 2 | Model 3 | Model 4 | Model 1 | Model 2 | Model 3 | Model 4 |
|  |  | GRS for educational attainment |  |  |  |  |  |  |  |
| Household income (£/year) | 276,779 | 1066 (<2e-16) | 1062 (<2e-16) | 874 (<2e-16) | 835 (<2e-16) | 1454 (<2e-16) | 1446 (<2e-16) | 1200 (<2e-16) | 1140 (<2e-16) |
| Body mass index (kg/m<sup>2</sup>) | 336,031 | -0.112 (<2e-16) | -0.111 (<2e-16) | -0.101 (<2e-16) | -0.097 (<2e-16) | -0.151 (<2e-16) | -0.150 (<2e-16) | -0.132 (<2e-16) | -0.129 (<2e-16) |
| Age at completion of full time education (years) | 228,886 | 0.0878 (<2e-16) | 0.0877 (< 2e-16) | 0.0844 (< 2e-16) | 0.0831 (< 2e-16) | 0.12 (<2e-16) | 0.119 (<2e-16) | 0.112 (<2e-16) | 0.109 (<2e-16) |
| Number of siblings (persons) | 332,037 | -0.0250 (<2e-16) | -0.0250 (<2e-16) | -0.0258 (< 2e-16) | -0.0253 (< 2e-16) | -0.038 (<2e-16) | -0.0382 (<2e-16) | -0.0293 (<2e-16) | -0.0279 (<2e-16) |
|  |  | GRS for height |  |  |  |  |  |  |  |
| Household income | 276,779 | 522 (<2e-16) | 515 (<2e-16) | 418 (1.8e-14) | 406 (2.7e-13) | 515 (<2e-16) | 509 (<2e-16) | 419 (1.7e-14) | 405 (2.9e-13) |
| Body mass index | 336,031 | -0.129 (<2e-16) | -0.128 (<2e-16) | -0.112 (<2e-16) | -0.116 (<2e-16) | -0.122 (<2e-16) | -0.121 (<2e-16) | -0.105 (<2e-16) | -0.109 (<2e-16) |
| Age at completion of full time education | 228,886 | 0.0350 (9.4e-09) | 0.0348 (1.1e-08) | 0.0289 (2.0e-06) | 0.0263 (2.0e-05) | 0.0349 (1.1e-08) | 0.0347 (1.2e-08) | 0.0286 (2.6e-06) | 0.0265 (1.8e-05) |
| Number of siblings | 332,037 | -0.0249 (<2e-16) | -0.0248 (< 2e-16) | -0.0130 (8.1e-06) | -0.0119 (7.2e-05) | -0.0264 (<2e-16) | -0.0263 (< 2e-16) | -0.0136 (3.0e-06) | -0.0127 (2.1e-05) |
|  |  | GRS for body mass index |  |  |  |  |  |  |  |
| Household income | 276,779 | -335 (1.8e-09) | -325 (5.2e-09) | -251 (4.0e-06) | -229 (3.4e-05) | -304 (4.7e-08) | -294 (1.3e-07) | -212 (0.00010) | -190 (0.0057) |
| Body mass index | 336,031 | 0.612 (<2e-16) | 0.611 (<2e-16) | 0.606 (<2e-16) | 0.606 (<2e-16) | 0.549 (<2e-16) | 0.547 (<2e-16) | 0.541 (<2e-16) | 0.541 (<2e-16) |
| Age at completion of full time education | 228,886 | -0.0219 (0.00032) | -0.0216 (0.00040) | -0.0201 (0.00092) | -0.0187 (0.0025) | -0.0231 (0.00016) | -0.0227 (0.00020) | -0.0201 (0.00096) | -0.0187 (0.0024) |
| Number of siblings | 332,037 | 0.0107 (0.00030) | 0.0105 (0.00036) | 0.00783 (0.0071) | 0.00750 (0.011) | 0.00130 (1.0e-05) | 0.00129 (1.3e-05) | 0.00850 (0.0035) | 0.00807 (0.0068) |

GRS = genetic risk score; PC = principal component; SD = standard deviation. The field contents are beta coefficients per 1 SD increase in GRS, with p-values for the linear association, testing the null hypothesis of no linear association between each observed trait and GRS in brackets. Statistical adjustment was performed as follows: model 1 – no adjustment; model 2 – adjustment for genotyping array only; model 3 – adjustment for genotyping array, 40 PCs and study participation centre; model 4 – adjustment for genotyping array, 40 PCs, study participation centre and non-linear regression terms for North and East axes of birth location.

As an alternative way to demonstrate the potential impact of such bias, we analysed simulated geographically-stratified complex traits which preserved coarse geographical variance in observed traits whilst removing direct genotype-phenotype effects. This analysis produced associations between GRS and complex traits even in the absence of direct genetic effects on biology, suggesting GRS predict geographical location within the UK Biobank sample (online methods and table S1).

The presence of structure within the genetic data of UK Biobank has several potential explanations, including a legacy of ancient ancestral groups that are not fully admixed^6,27^, a consequence of non-random mating or polygenic selection^28–30^, a study artefact induced by selection bias^17^ or a combination of all these explanations. Regardless of origin, unaddressed structure in this sample is sufficient to mean that predictions based on GRS are capable of inducing associations where there is little or no direct effect. Recent evidence from an investigation in the USA^31^ also illustrates associations between GRS and complex traits at the ecological level. Now manifest, this property should be added to the growing list of limitations to naïve use of GRS - including horizontal pleiotropy^7^, high false discovery rate^32^, association with coarse ancestral groups^33^ and prediction of inter-generational phenotypes which complicates interpretation^34^.

The ability of very large studies to detect effects indistinguishable from artefactual biases or ancestral differences demands reworked approaches to exploit^35^, or at least account for, structure. Exciting recent developments aim to improve statistical models^36^ or leverage information from family-based study designs for unbiased inference^37^. Until such methods have developed further, the truth is that a thorough understanding of the properties of genotypic and phenotypic data and impact of study design will remain critical in allowing reasonable inference.

## Author Contributions

NT, SH and ND conceived the study; SH, DL, RM and ND performed the analysis; SH, DL and NJT wrote the paper. All authors discussed the result and commented on the paper.

## Acknowledgements and funding

We are extremely grateful to all the families who took part in this study, the midwives for their help in recruiting them, and the whole ALSPAC team, which includes interviewers, computer and laboratory technicians, clerical workers, research scientists, volunteers, managers, receptionists and nurses.

The UK Medical Research Council and Wellcome (Grant ref: 102215/2/13/2) and the University of Bristol provide core support for ALSPAC. A comprehensive list of grants funding is available on the ALSPAC website at (http://www.bristol.ac.uk/alspac/external/documents/grant-acknowledgements.pdf).

The UK Medical Research Council (MRC) and the University of Bristol support the MRC IEU. NJT is a Wellcome Trust Investigator (202802/Z/16/Z), a work-package lead in the Integrative Cancer Epidemiology Programme (ICEP) that is supported by a Cancer Research UK programme grant (C18281/A19169) and works within the University of Bristol NIHR Biomedical Research Centre (BRC). DL is funded by Wellcome Trust and Royal Society (Grant ref: WT104125MA). The Economics and Social Research Council (ESRC) support ND via a Future Research Leaders grant (Grant ref: ES/N000757/1). GDS is the director of and a programme lead in the MRC-IEU (MC_UU_12013/1). GH receives funding from the Wellcome Trust (Grant Ref: 208806/Z/17/Z). KHW is funded by programmes 3 and 4 of the MRC IEU (Grant refs; MC_UU_12013/3 and MC_UU_12013/4), and by Wellcome Trust funding (Grant ref: 202802/Z/16/Z awarded to NJT). SH receives support from Wellcome (Grant ref: 201237/Z/16/Z). No funding body has influenced data analysis or interpretation. This work was carried out using the computation facilities of the Advanced Computing Research Centre-http://www.bris.ac.uk/acrc/ and the Research Data Storage Facility of the University of Bristol-http://www.bris.ac.uk/acrc/storage/. This research was conducted using the UK Biobank Resource applications 8786 and 15825.

We wish to acknowledge the contributions of Professor Augustine Kong of the Big Data Institute, Oxford University. Professor Kong helped in the preparation of this manuscript through discussion and development of themes central this work.

This work arose from discussion within the MRC IEU dry lab meeting group, which is a community of users of genetic data at the MRC IEU. This group meets regularly to discuss analysis of genetic data and observations during these meetings formed the starting point for this work. We are very grateful to all the members of this group for their input.

